# Fossils matter: improved estimates of divergence times in *Pinus* reveal older diversification

**DOI:** 10.1101/073312

**Authors:** Bianca Saladin, Andrew B. Leslie, Rafael O. Wüeest, Glenn Litsios, Elena Conti, Nicolas Salamin, Niklaus E. Zimmermann

## Abstract

**Background:** The taxonomy of the genus *Pinus* is widely accepted and a well-resolved phylogeny based on entire plastome sequences exists. However, there is a large discrepancy in estimated divergence times of major pine clades among existing studies mainly due to differences in fossil placement and dating methods used. We currently lack a dated molecular pine phylogeny that makes full usage of the rich fossil record in pines. This study is the first to estimate the divergence dates of pines based on a large number of fossils (21) evenly distributed across all major clades in combination with applying the most novel dating method.

**Results:** We present a range of molecular phylogenetic trees of *Pinus* generated within a Bayesian framework using both the novel fossilized birth-death and the traditional node dating method with different fossil sets. We find the origin of pines likely to be up to 30 Myr older (Early Cretaceous) than inferred in most previous studies (Late Cretaceous) and propose generally older divergence times for major clades within *Pinus* than previously thought. Our age estimates vary significantly between the different dating approaches but the results generally agree on older divergence times. We present a revised list of 21 fossils that are suitable to use in dating or comparative analyses of pines.

**Conclusions:** An accurate timescale for the divergence times in pines is essential if we are to link diversification processes and functional adaptation of this genus to geological events or to changing climates. Next to older divergence times in *Pinus*, our results indicate that node age estimates in pines depend on dating approaches and fossil sets used due to different inherent characteristics of dating approaches. Our set of dated phylogenetic trees of pines presented herein provide the basis to account for uncertainties in age estimations when applying comparative phylogenetic methods, which will improve our understanding of the evolutionary and ecological history in pines.

## Background

The genus *Pinus*, with approximately 115 extant species, is the largest genus of conifers and one of the most widely distributed tree genera in the Northern Hemisphere [1]. Pines are an integral component of many Northern Hemisphere ecosystems, and they have a well-documented, rich fossil record [2] stretching back as much as 130 - 140 million years [3, 4]. Many studies have focused on this genus, particularly with regard to its phylogenetic relationships [1, 5-10], ecology [11, 12], biogeography [13, 14], and the timing of diversification events [15]. There exists a wealth of molecular, morphological and fossil data on the genus. However, no study has yet made full use of all existing data to generate both a fully resolved phylogenetic tree including all extant species and a time calibration of such a tree based on the rich fossil record. Such extensively dated and comprehensive phylogenetic trees will allow us to fill significant gaps in our understanding of the evolutionary and ecological history in pines [16].

The genus *Pinus* has traditionally been divided into two major clades based on the number of vascular leaf bundles (either one or two bundles, corresponding to sections *Haploxylon* and *Diploxylon* respectively, also referred to as subgenera *Strobus* and *Pinus*) [1]. Previous studies have not been able to consistently resolve relationships within these major clades, resulting in a number of different sectional and subsectional classifications. In 2005, Gernandt *et al*. [5] proposed a new classification based on phylogenetic trees inferred from two chloroplast genes, dividing the pines into two subgenera (*Pinus* and *Strobus*), four sections (sections *Pinus* and *Trifoliae* in subgenus *Pinus* and sections *Parrya* and *Quinquefoliae* in subgenus *Strobus*) and 11 subsections (*Australes, Ponderosae, Contortae, Pinus, Pinaster, Strobus, Krempfianae, Cembroides, Balfourianae* and *Nelsoniae*). Although taxonomically comprehensive and widely accepted, their study relied exclusively on sequences from the *matK* and *rbcL* genes, and was thus unable to resolve relationships within several of the subsections. Subsequent studies have improved phylogenetic resolution, but have mostly focused on specific subclades [e.g. 9, 13, 17, 18]. More recently, Parks *et al.* [6] analyzed the entire chloroplast genome for 107 pine species, which largely confirmed the structure proposed by Gernandt et al. [6] and provided better resolution for much of the tree. However, despite the detailed chloroplast data and the availability of potential fossil calibration points, comprehensive time-calibrated molecular phylogenetic trees are lacking.

Sound estimations of divergence times within phylogenetic trees benefit from using many fossils that are evenly distributed across the tree, a strategy that better accounts for rate variation when using relaxed molecular clock models [19-21]. In addition, multiple calibrations can overcome negative effects from errors in dating and placement of single fossils [22]. In the genus *Pinus*, a rich fossil record exists, with the first fossil appearing in the Early Cretaceous [3, 4]. Beside Mesozoic pine fossils [3, 4, 23-26], numerous fossils were described from the Cenozoic era and placed within various pine clades [27-32]. However, most of the recent time calibrations of pine divergences have typically used very few (usually 1-3) fossils [11, 13, 15, 16, 18] (but see [14]). Some of these fossils are controversial regarding their phylogenetic assignment and age (e.g. the use of *P. belgica* as discussed in [15]), leading to inconsistent age estimates of the origin of pines and divergence times of subsections therein. There remains a great need to include a larger number of carefully evaluated fossil constraints, preferably evenly distributed across all major clades, to improve our understanding of pine evolution.

Although Bayesian methods using a relaxed molecular clock are widely accepted for time calibration of molecular trees, there is ongoing debate regarding the best strategy to convert fossil information into calibration information [33-36] and methods are still under development [34]. In the widely and traditionally used *node dating* method [termed by 36] (ND, hereafter), the geological age of the oldest fossil of a specific clade is transformed into a calibration density (also referred to as prior for divergence times [37] or probabilistic calibration priors [38]) to assign a known age range to the stem node (also referred to as calibrating nodes [39]) of the respective clade in the phylogenetic tree [34]. The probabilistic calibration prior accounts for uncertainties underlying the age of the fossil and the likelihood that the true divergence occurred before its first appearance in the fossil record [19, 37]. However, there is no objective way to define the calibration densities and researchers have used different approaches to define them [19, 37, 38, 40]. Recently, the *fossilized birth-death* (FBD, hereafter) method has been introduced as a new approach for time calibration of molecular phylogenetic trees [41, 42]. This method assumes that extant species and fossils are both part of the same evolutionary process and it does not constrain fossils to nodes only. Rather fossils represent extinct tips and are placed along specific branches of the phylogenetic tree anywhere within the lineage a fossil is assigned to following the specific birth-death process estimated. This allows including all fossils as tips within a clade instead of summarizing them into calibration densities assigned to nodes as in ND. The FBD method therefore overcomes some of the known shortcomings of the ND method (well discussed in the literature [36, 42]) and is considered promising [16]. While FBD has the potential to be widely used in the future [43], only few studies have directly compared these two time dating methods so far [44, 45]. No conclusion has been reached to date as to whether estimated divergence times are in agreement between the two methods [44-46].

Here, we build for the first time a comprehensive phylogenetic tree for all described species of *Pinus* that is dated with a large number of fossils evenly distributed across all major clades using the novel FBD method. More specifically, our objectives are: (1) to provide a revised and well-supported time-scale for the evolution of major subsections of pines; (2) to test the sensitivity of age estimates to different dating methods and fossil sets; and (3) to provide a revised list of fossils and their phylogenetic placement within the genus for use in further studies on pine evolution.

To achieve these goals we infer phylogenetic trees based on eight chloroplast sequences within a Bayesian relaxed molecular clock framework using both the novel FBD and, for comparison, the traditional ND method (Figure 1). In ND we apply two different prior approaches for assigning calibration densities on nodes. The first approach follows what was applied in previous pine studies [11, 13, 15] and is based on calibration densities that reflect the geological timescale of the layers in which the fossils were excavated. As we believe that this approach may define too tight calibration densities on nodes that do not reflect the uncertainty in our prior knowledge (especially toward older nodes), we also defined an alternative approach. In this second approach, we constructed calibration densities of increasingly higher uncertainty with increasing age, which better accounts for uncertainty in the *a priori* information of calibration constraints. In both methods (FBD and ND) we estimated the absolute age scale of the phylogenetic trees from two sets of fossils for each setting (14 or 21 fossils, resp. 12 and 15 in ND due to using only the oldest fossil per node). The two fossil sets differ in our confidence regarding fossil ages and phylogenetic assignments. Our study therefore provides improved estimates of divergence times in pines.

**Fig 1.**
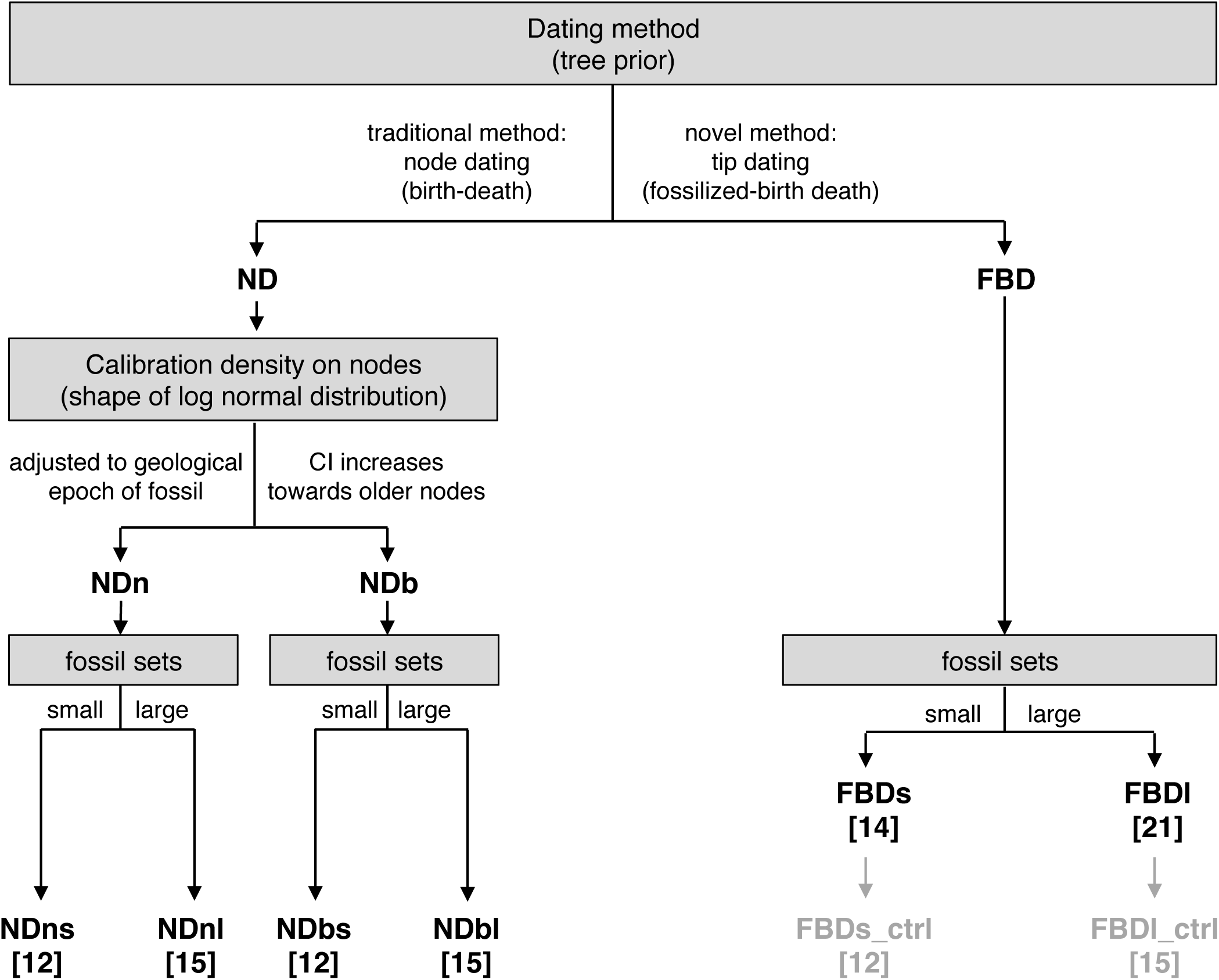
Flow chart illustrating the different dating methods applied. We used both the node dating (ND) and the fossilized birth-death (FBD) method. In ND, we defined the calibration densities on calibration nodes either with narrower (NDn) or broader (NDb) density distributions of priors on age. Each dating was carried out with a smaller (s) or larger (l) fossil set (fossil number for each approach indicated in square brackets). Since node dating only uses the oldest fossil per node, this resulted in fewer fossils used in the small and the large fossil set in ND compared to FBD. A control run (FBDs/l_ctrl) was additionally executed for FBD in which exactly the same fossils as in NDs/l were used.

## Results

### Divergence times in *Pinus*

Our analyses suggest a *Pinus* crown lineage diversification likely in the Early Cretaceous based on the FBD method (node *a* in Figure 2), irrespective of using the small fossil set (FBDs: median: 125 Ma, 95% confidence interval, CI: 144 - 106 Ma; Figure 3) or the large fossil set (FBDl: median: 124 Ma, 95% CI: 145 - 105 Ma). The estimates from the ND method defined with broader calibration densities on nodes (NDb) support an Early Cretaceous age using both the small and large fossil sets, although slightly younger than FBD (NDbs: median: 112 Ma, 95% CI: 156 Ma - 96 Ma; NDbl: median: 110 Ma, 95% CI: 150 Ma - 96 Ma). In contrast, the ND approach based on the geological time scale defined with narrower calibration densities (NDn) estimates significantly younger, Late Cretaceous divergence ages using both fossil sets (NDns and NDnl: median: 90 Ma, 95% CI: 96 - 90 Ma). In FBD, crown splits within the two subgenera (node *b* and *c* in Figure 2 and 3) are estimated to have diverged at roughly the same time during the Late Cretaceous to early in the Paleocene (FBDs: median subgenus *Pinus*: 64 Ma, 95% CI: 87 - 52 Ma; median subgenus *Strobus*: 68 Ma, 95% CI: 89 - 53 Ma; FBDl: median subgenus *Pinus* 69 Ma, 95% CI: 92 - 56 Ma; subgenus *Strobus* median: 71 Ma, 95% CI; 92 - 56 Ma). In the ND method, these two splits are estimated to have diverged in the late Paleocene and early Eocene. Figure 3B illustrates the crown age estimates of sectional and subsectional nodes for the methods and approaches used. Maximum clade credibility trees (MCT trees) of all dating methods, approaches and fossil sets are provided in the Additional file 1. Most node age estimates in previous studies revealed younger ages than our FBD and NDb and partly also than our NDn methods (Figure 3B).

**Fig 2.**
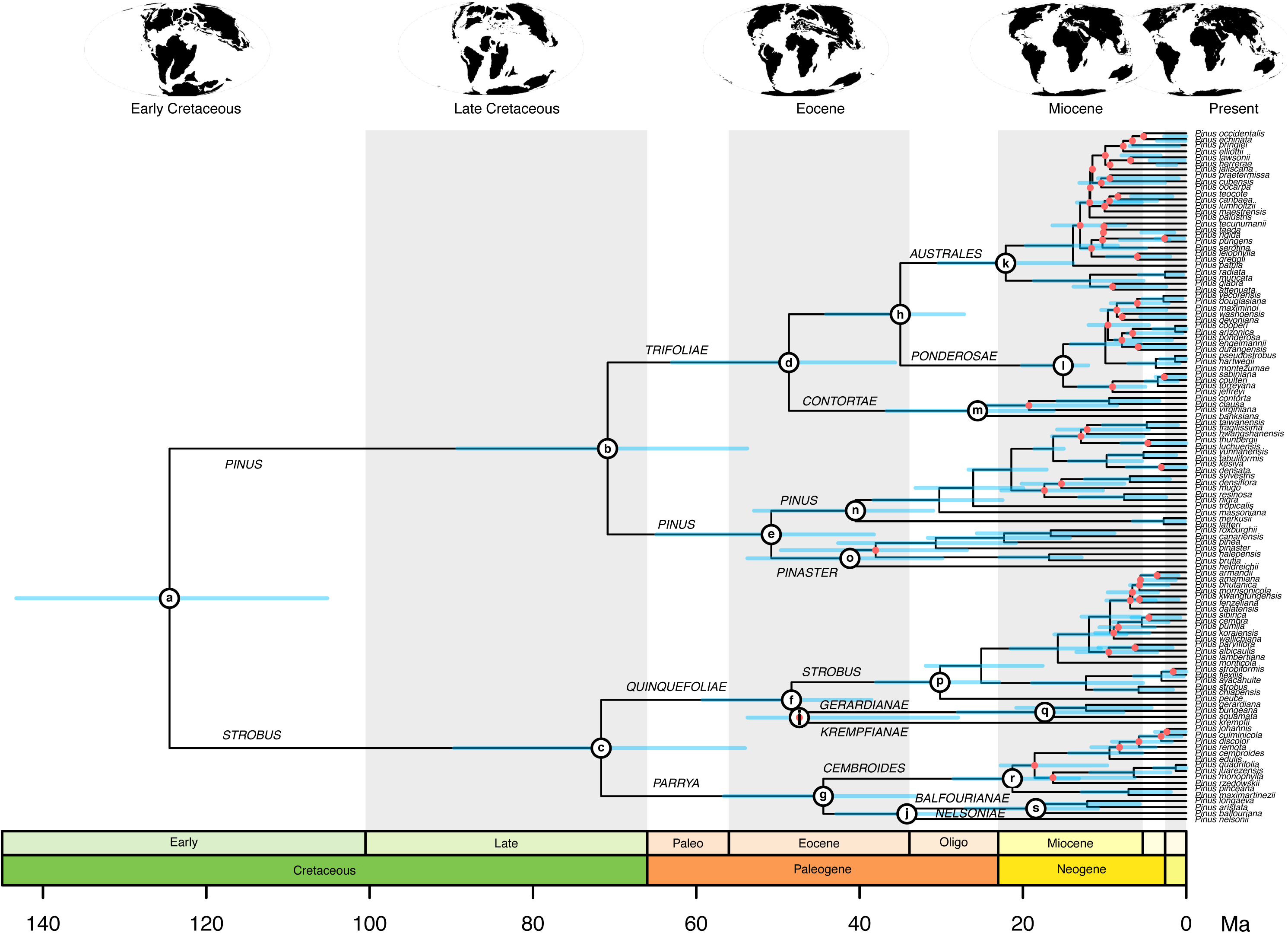
Inferred maximum clade credibility (MCC) tree from results of the FBDl method (fossilized birth-death method, based on the larger set of 21 fossils). Nodes with red dots indicate Bayesian posterior probabilities lower than 0.95, while all other nodes have posterior probabilities higher than 0.95. Light blue lines on nodes represent the 95% highest posterior density (HPD) of the inferred phylogenetic trees. The node labels (*a-s*) indicate those nodes represented in Figure 3. The geological timescale is in million years and the paleogeographic maps on top were redrawn from [77].

**Fig 3.**
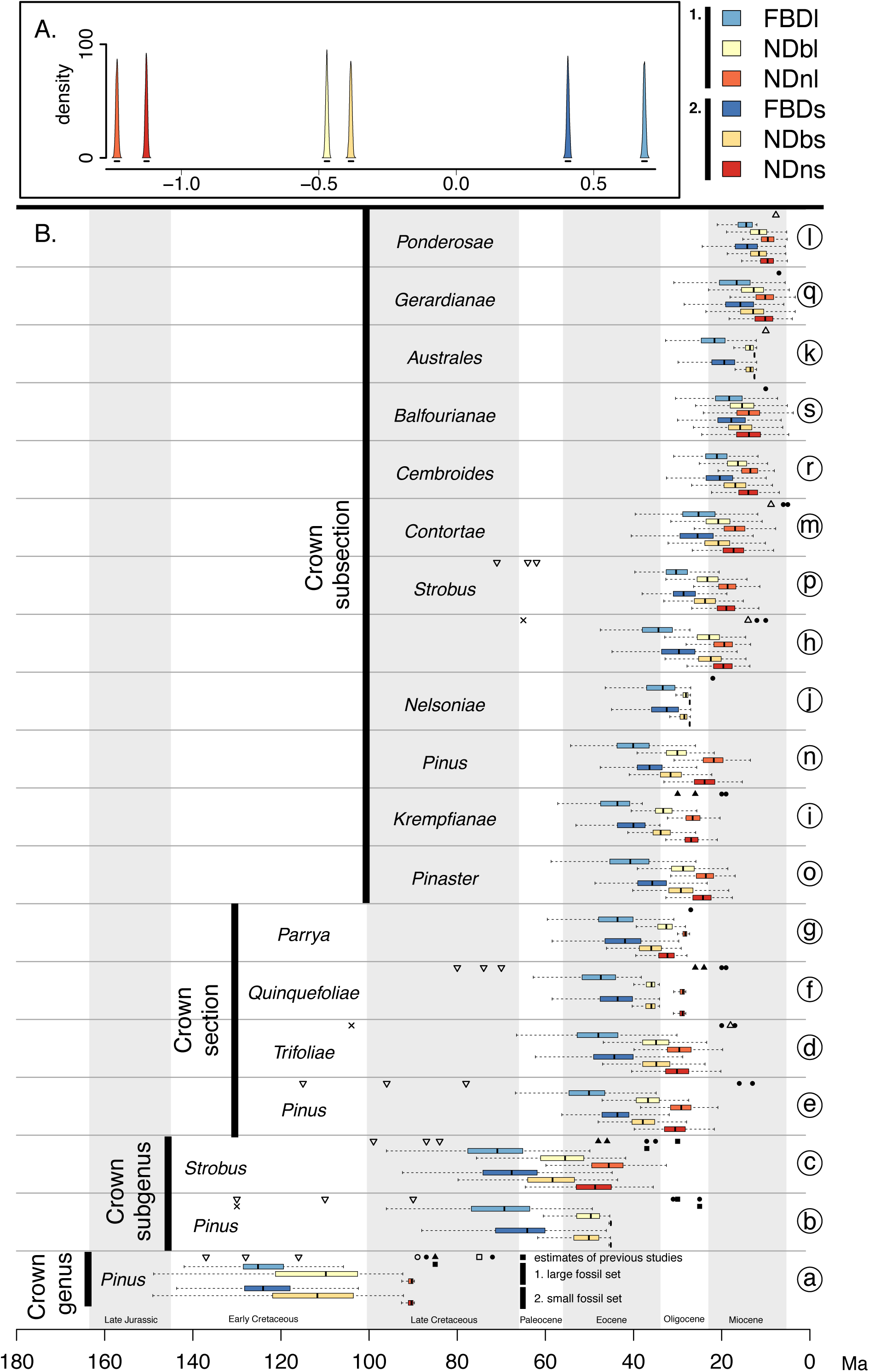
Comparison of estimated node ages of the 19 major clades of *Pinus* across all applied dating approaches. **A**: The density of effect sizes originate from a mixed-effect model and illustrate to what degree the estimated node ages differ among dating approaches (different colors; see legend) and among fossil sets (1. darker colors for the large, 2. brighter colors for the small fossil set; see legend). The 95% confidence intervals of effect sizes are illustrated with a line below the density curves. Non-overlap of these intervals indicates significant difference on node ages among all 19 nodes. **B**: Boxplots illustrate the estimated node ages across dating approaches and fossil sets for the major clades (*a-s* illustrated in Figure 2). Whiskers span the 95% highest probability density (HPD), while boxes span the 50% HPD, with the median node age indicated by a vertical bar. The x-axis indicates the geological time in million years. Symbols represent average node ages as estimated in the following studies: Gernandt *et al.* [16] (filled circle), illustrating estimates resulting from two different calibration scenarios; Hao *et al.* [13] (filled upward triangles); Willyard *et al.* [15] (filled squares), illustrating the estimates based on both the chloroplast and the nuclear sequence data, but only presenting results of their 85 Ma calibration scenario as this was indicated to be more realistic; Hernandez-Leon *et al.* [18] (open upward triangle); He *et al.* [11] (open circles); Leslie *et al.* [47] (open squares); Geada Lopez [48] (crosses); Eckert *et al.* [14] (open downward triangle). The following abbreviations are used. FBD: fossilized birth-death method; ND: node dating method; l: analyses based on the large fossil set; s: analyses based on the small fossil set; n: narrow calibration priors in ND based on the geological age of the respective fossil; b: broad calibration priors in ND.

### Comparison of dating methods

In general, the different methods produce several consistent patterns among the 19 nodes representing crown nodes of subsection and higher-level clades (Figure 3, nodes *a*-*s*). First, the FBD method estimates significantly older ages for the 19 nodes than the ND method, irrespective of the specific calibration employed or fossil set used (Figure 3A). Second, NDn (narrower distribution of calibration densities) estimates significantly younger ages for the 19 nodes in the phylogenetic trees than does NDb (broader distribution, Figure 3A), particularly for the crown age (Figure 3B). Third, FBDl analysis (21 fossils) provide significantly older estimates for the 19 nodes than do FBDs analysis (14 fossils, Figure 3A). In contrast, in both ND methods, applying the large set of 15 fossils lead to slightly but significantly younger age estimates than do those based on the small set of 12 fossils (Figure 3A). Control runs of FBD using the same 12 and 15 fossils as in ND reveal very similar node ages as in FBDs (14 fossils) and FBDl (21 fossils) except at the crown node of *Pinus* (Additional file 2).

### Sensitivity of dating methods to prior settings

We examine the relative influence of the probabilistic calibration priors and the sequence data on the Bayesian age estimates in each method (Figure 4) by comparing the effective prior distributions to the posterior distributions of the calibration nodes. We find significant differences across dating methods: the calibration priors in the NDn method are highly similar to the posterior age estimates, revealing the strong influence of the defined calibration priors on estimated node ages. In contrast, the NDb and FBD approaches reveal increasingly lower influences of the calibration priors, indicating a lower sensitivity of posterior age estimates to calibration priors. This pattern emerges irrespective of the fossil set used (Figures 4A and 4B).

**Fig 4.**
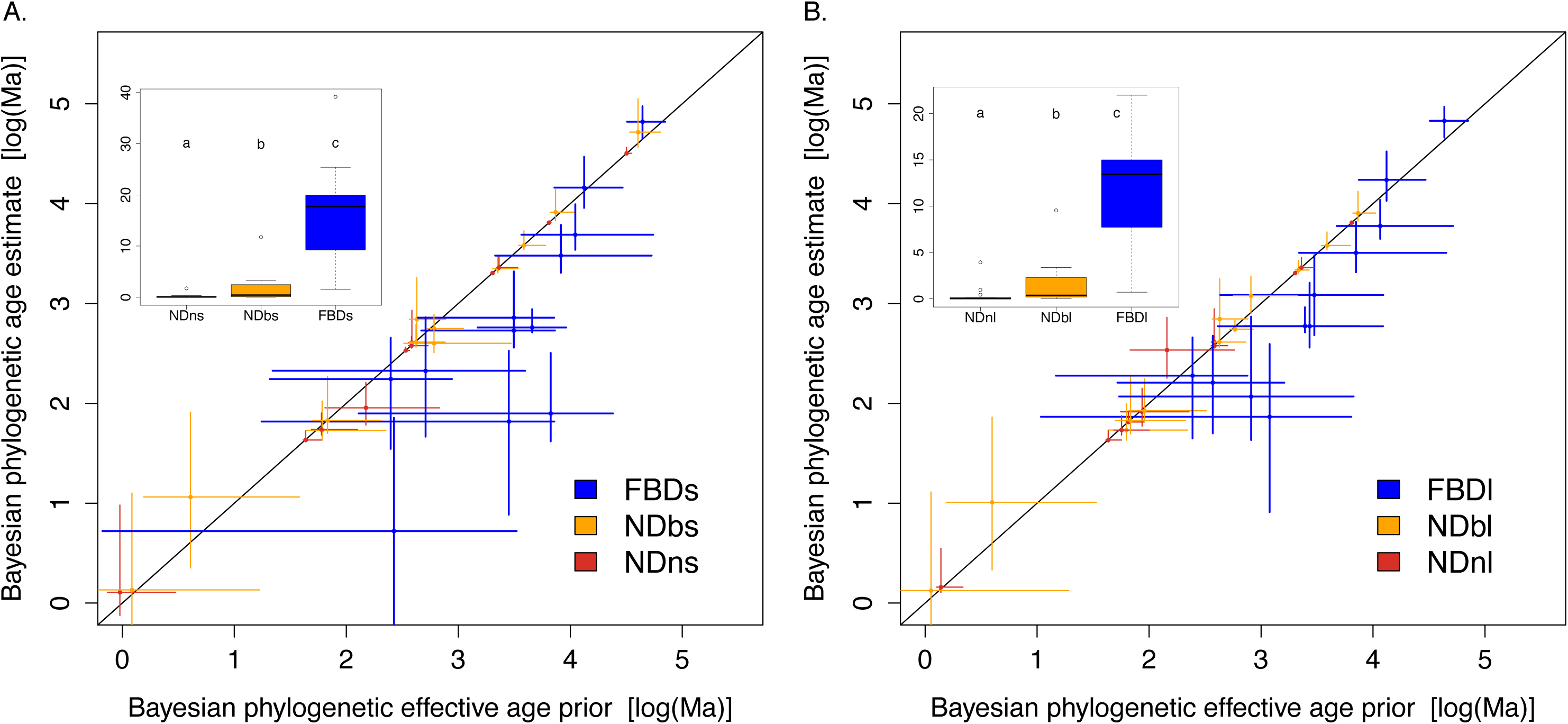
Comparison of the effective age prior density against the posterior calibration densities (Bayesian phylogenetic age estimate) for the three dating approaches used (FBD: fossilized birth-death (blue); NDn/b: node dating with narrow (red) and broad (orange) prior distributions; s/l: small (**A**) and large (**B**) fossil sets. Boxplots represent the absolute deviation from the 1:1 line, while letters indicate significant differences in absolute deviations at the level of p = 0.05 (based on a paired Wilcox test).

### Sensitivity of node age estimates to single fossil exclusions

The exclusion of a single fossil can have a strong influence on the calibration of single node ages. Figure 5 illustrates how much the 19 nodes (*a-s*) differ in calibrated ages when leaving out the individual fossils in FBDs (Figure 5A) and FBDl (Figure 5B). See Additional file 3 for this same sensitivity analysis with the ND-based phylogenetic trees. The two fossils *P. fujii* and *P. crossii* lead to generally younger node ages on almost all 19 nodes when left out, both in FBDs and FBDl (Figure 5) and in NDb approaches (Additional file 3). In NDn approaches, node ages are not sensitive to the exclusion of *P. fujii* but they are sensitive to inclusion of *P. halepensis* and *P. crossii*. In contrast, excluding the oldest fossil (*P. yorkshirensis* in FBD and *P. triphylla* in ND) leads to generally older ages, especially in older nodes. In FBD the same is observed for *P. haboroensis*, while in NDb the fossil *P. baileyi* also leads to older ages when excluded (Additional file 3). The exclusion of all other fossils does not have a strong effect on the ages of the19 nodes.

**Fig 5.**
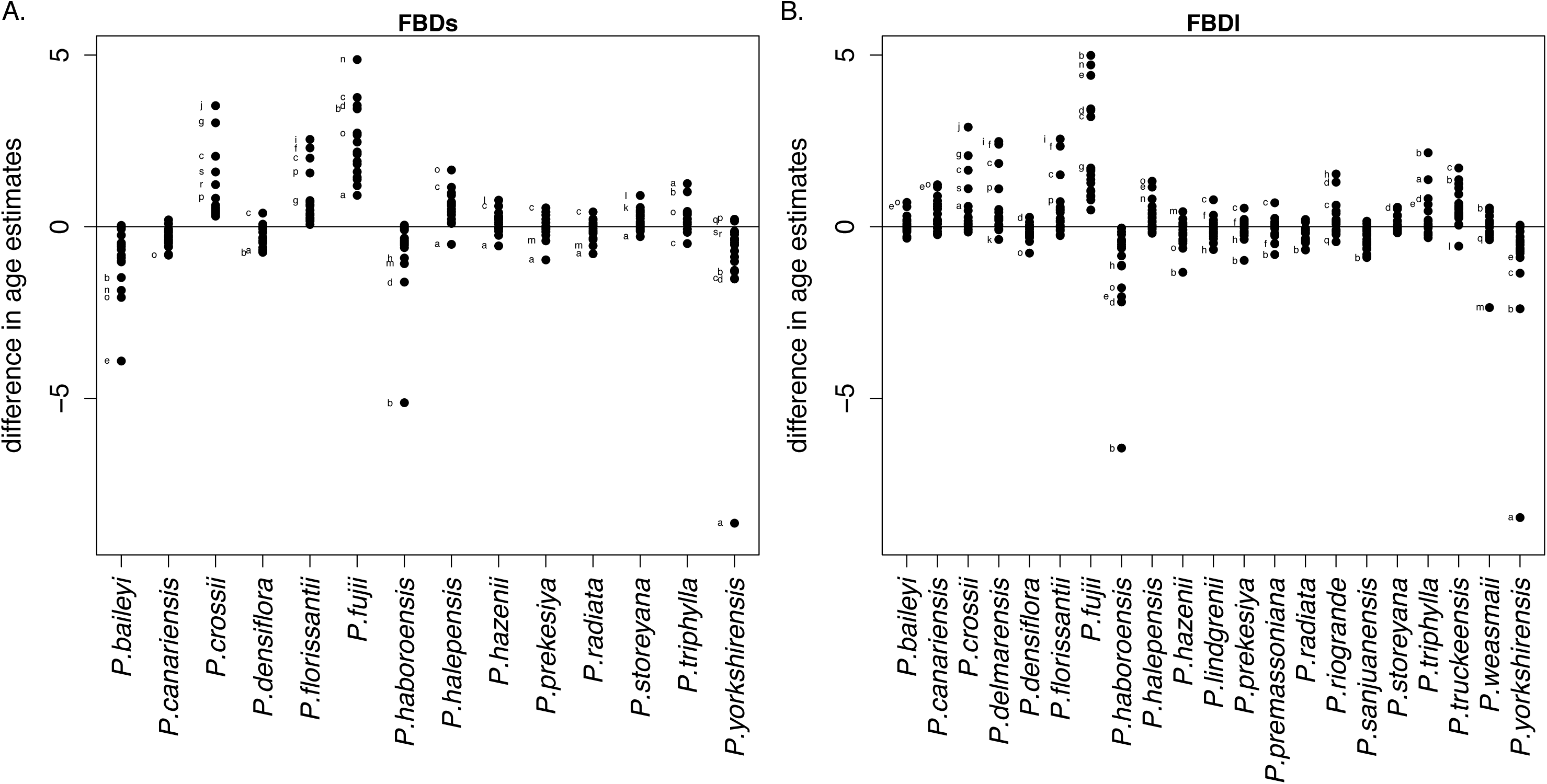
Sensitivity of the time calibration to single fossil exclusion for the fossilized birth-death approaches (FBD). This test measures the difference in age estimates of the 19 major nodes (*a-s*, see also Figure 3) when keeping versus removing single calibration constraints (fossil, labeled on x-axis) at a time. Illustrated are the results from using the small (**A**) and the large (**B**) fossil set. Letters (see figure 2 for assignment) indicate nodes with highest deviations.

## Discussion

### Divergence times in *Pinus*

The Late Cretaceous crown age of *Pinus* inferred in our study (FBD and NDb) is approximately 30 Myr older than the age estimated in most previous studies [13, 15, 16, 47, 48] (Figure 3B). To our knowledge, only one other study estimated a similarly old crown age in *Pinus* [14] using the fossil *P. belgica* [23] whose exact phylogenetic assignment and age are uncertain [15, 47]. For this reason we did not use it in our study. Our estimated crown age is consistent with the very recent discovery of the now oldest fossil attributed to the genus *Pinus* (*P. mundayi* sp.), which has been dated to the Early Cretaceous (Valanginian, ca. 133-140 Ma) [4] but was not included in our study due to its disputed placement [49]. Although our genus crown-age estimates are similar to the one found in the study using *P. belgica*, the crown ages of subgenera and sections inferred in our study are clearly younger than their estimate [14] (Figure 3B).

In line with the early crown age of the genus *Pinus*, we also found strong evidence for a Late Cretaceous to Paleocene origin of the crown of the two *Pinus* subgenera (supported by most dating methods), which clearly differs from the Eocene origin suggested in most previous studies [13, 15, 16, 47]. Further, most subsections were thought to have emerged during the Miocene [15], but our results support this conclusion only in half of the subsections (*Ponderosae, Gerardianae, Australes, Balfourianae, Cembroides*), while the other subsections date back to the Oligocene and the Eocene. Still, irrespective of the estimated subsectional crown ages, the major diversification of extant species occurred during the Miocene.

### Uncertainties in divergence time estimations

Divergence time estimates were found to be dependent on the number of fossils used [39], dating methods [44-46], phylogenetic assignment of fossils [14] and uncertainty of fossil ages [15]. Several reasons are likely responsible for the large discrepancies in estimated node ages among the different studies previously published on pines, including our own results. In the following, we discuss the likely reasons for these differences.

#### Effect of fossil numbers

Our study used more fossils than previous ones, and this is likely one reason for the difference in divergence estimates. Dating methods based on only few fossils are very sensitive to the fossil assignment and defined calibration priors and can lead to biased substitution rate estimates [39]. If the prior on divergence time derived from a single fossil is inaccurate, then the estimated ages of all nodes are affected, because there are no other calibration points that can ameliorate the effects from this error [22, 50]. Even if the single used fossil is accurately placed in the phylogeny, age estimates of nodes distant from the calibration point may still be prone to inaccuracies [35]. Using multiple calibrations evenly distributed across all major clades is crucial to inform the relaxed molecular clock and model rate variation among lineages [20, 22]. Multiple calibration constraints therefore lead to more precise and robust dating analyses [39]. The greater number and more even distribution of fossils across the phylogeny used in this study is an important step towards an accurate calibration of the molecular clock within *Pinus*.

#### Effect of dating method

Node age estimates not only depend on the number of fossils used for calibration of the molecular clock, but clearly also on their placement, and the method used for calibration (Figure 3). Our study is the first in pines to have used FBD. The reason for observing significantly older age estimates in FBD than in ND is primarily due to differences in the placement of fossils between FBD and ND and not due to the larger fossil numbers used in FBD compared to ND. Indeed, FBD runs with exactly the same 12 or 15 fossils as in ND (FBD_ctrl) revealed node ages very similar to those inferred by using the 14 or 21 fossils (Additional file 2). In ND, we placed fossils at nodes, while in FBD we let the same fossil be placed anywhere along the (stem) branch or even anywhere within the clade. This pushes the divergence time of the crown node of the respective clade deeper in time in FBD than in ND. Another major difference between FBD and ND is that calibration points are directly used in the stochastic process underlying the FBD approach, which leads to a better distribution of branching times from the birth-death process. In contrast, ND discards the branching times that do not correspond to the calibrations by simply multiplying the tree prior density with the calibration densities [51]. Taking the product of the two densities does not produce a proper conditional birth-death sampling prior, in contrast to the FBD method, and this approach can also lead to bias.

We assigned fossils to clades based on our paleontological knowledge of traits. FBD randomly places fossils along a specified branch or within this clade. This random assignment could be further improved (and better constrained) by using morphological data matrices for extant and dated fossil taxa. The novel dating methods (including FBD or total-evidence dating, TED [36]), which were developed to jointly analyze morphological matrices of extant and extinct species and molecular data under models of morphological character evolution, are attractive and promising [52, 53]. However, these methods are still contentious as they are actively being developed and debated [33, 53, 54]. Previous analyses based on the TED method in combination with morphological data have consistently led to older divergence age estimates [reviewed in 54] and have consequently been criticized as not yet being fully developed. For *Pinaceae*, Gernandt et al. [16] were the first to demonstrate that the combination of morphological and molecular data can improve the accuracy of divergence time estimates and phylogenetic relationships. Additional morphological matrices for fossil and extant Pinaceae, including some pines, are available [55, 56], although not yet for sufficiently many pine species and could therefore not be used in this study. This is a demanding task as it requires a reevaluation of many described pine fossils in relation to a list of diagnostic features of modern clades, but also more detailed anatomical descriptions of most living species. Such anatomical descriptions would benefit from making full use of the 21st century technology (e.g. high resolution computed tomography scanning [57]).

The few existing analyses based on FBD did not reveal a general trend towards over or underestimation of node ages. One study on tetraodontiform fishes had inferred similar ages using ND and FBD methods [44], while another study on penguins inferred much younger age estimates with FBD combined with morphological characters than found in previous studies using node dating [41]. However, despite identified drawbacks in FBD (as discussed in [33, 53]), this method clearly overcomes major shortcomings of ND, most importantly the arbitrariness underlying the assignment of probabilistic calibration priors.

Age estimates of ND are sensitive to the defined probabilistic calibration priors [19, 58, 59]. No objective approach has yet been suggested to define the priors for divergence times and it has been shown that incorrect calibration constraints negatively affect divergence estimates [19, 60]. This sensitivity to calibration priors is also visible in our study. First, we found significant differences in the age estimates between NDn and NDb where different density shapes were used (Figure 3A), indicating that the derived posterior node ages are strongly influenced by the assumed prior calibration densities. Second, the prior sensitivity analyses (Figure 4) revealed that posterior age estimates in NDn are significantly more sensitive to the effective priors of calibration constraints than in NDb, whereas FDB is least sensitive. This likely affected the Bayesian analyses and may have led to biased age estimates. The broader calibration densities in NDb have led to age estimates that were less strongly influenced by prior settings on calibration densities. Unless one is certain about narrow prior densities, it seems more conservative to define them broadly and allow for a more balanced influence of both the molecular data and the priors of calibration densities. However, there remains the trade-off between defining too narrow prior densities (that can bias the estimation) and too broad prior densities (that will lead to overly large uncertainties). The molecular data itself has very little information to estimate the relaxed clock. The information is coming from the calibration constraints, and these will not be useful if defined too broadly. In our study, we have more confidence in the age estimates of NDb than NDn, which are consistent with the older divergence times for pines compared to previous studies. Another known shortcoming in ND is the difficulty to specify multiple node calibrations, especially when one node is ancestral to the other [53], which often occurs when many fossils are used within a clade. As the priors of these multiple constraints interact, the effective prior distributions may be quite different from the initially set prior distributions that were defined based on biological and paleontological knowledge [51]. This is also the case in our study (Additional file 4), where some effective prior distributions of ages were slightly changed, mainly truncated and sometimes slightly shifted compared to the initially set priors (examples of such fossils: *P. premassoniana*, *P. densiflora*, *P. storeyana* in Additional file 4). One of the biggest differences can be found for the prior on the calibration node of the fossil *P. premassoniana*. This could be due to the long branch of extant *P. massoniana* that had most probably already emerged during the Oligocene, while the assigned fossil age is comparably younger. The older posterior distribution indicates that most probably an older age should be assigned to this calibration node, because the fossil likely does not represent the oldest possible age for this node. Another example where the specified and effective priors differ is in the case of the calibration constraint based on *P. storeyana*. Here, it is possible that this fossil should be placed in the crown group of the “attenuatae-group” within subsection *Australes* (see discussion Additional file 7A).

In summary, our age estimates vary between the different methods and fossil sets used, due to the discussed inherent characteristics of each, but the pattern of FBD and NDb, in which we have confidence, is overall very consistent, also compared to the large range of age estimates available from previous pine studies.

#### Which fossil set?

Large sensitivities in age estimation to the exclusion of single fossils (Figure 5) may occur (i) if a fossil has a strong influence on node age estimates, (ii) if its phylogenetic placement is wrong, or (iii) if its assigned age is incorrect. A strong influence (point i) may arise if no other phylogenetically nearby fossil contributes to support the age estimate around the one that has been removed. The oldest fossil of the set has a strong influence, but such effects can also occur among younger fossils (Figure 5). Certainly, some calibration points are more reliable (points ii and iii) than others [19], and adding too many unreliable fossils may also bias estimates of rates and dates. The larger fossil set in our study includes (in addition to the fossils of the smaller set) fossils in which we have somewhat less confidence regarding phylogenetic assignment and age. If the addition of the seven “riskier” fossils to the larger fossil set had added considerable bias to age estimates, one could expect that the estimated ages would change noticeably when removing these fossils from the dating analyses. We would also expect that the effect would be more severe than when leaving out one of the 14 “conservative” fossils. Including those additional seven fossils led to significantly older (in FBD) or significantly younger (in ND) ages, but the differences were consistent across the methods and fairly small. More importantly, the fossils to which the node age estimates were most sensitive to when single fossils were left out (Figure 5) did not include any of those seven “riskier” ones. We conclude that our age estimates are robust towards changes in the fossil set, and that the distribution of fossils across the phylogenetic tree seems to be defensible, as we find similar results regardless of the fossil set used. We therefore suggest using the tree calibrated from the larger fossil set, which allows for the inclusion of all available information to calibrate the relaxed clock models for improved divergence time estimation in pines.

### Implications and conclusions of revised diversification times in *Pinus*

Pines are exceptionally interesting for studying the link between the evolution of physiological adaptations, functional traits and ecological niches (as e.g. [11, 61-65]). They have adapted to almost every forest habitat on the Northern Hemisphere while exhibiting a large range of observed morphological and physiological traits. The generally older divergence times found in our study compared to previous studies have consequences for our understanding of the evolution and biogeographic history of *Pinus*, because splits among important clades have to be interpreted within a climatic and tectonic context (Figure 2). For example, corridors for high latitude migration became increasingly reduced as the Atlantic Ocean widened and the climate started to fluctuate over the Cenozoic [2], which may have affected the origin and diversification of major clades. A sound understanding of the biogeographic history of pines, and its relation to climatic drivers and geographic constraints, requires accurately dated divergence times. This is particularly true for the major crown clades that diversified over the Cenozoic, and whose current diversity most likely relates to major climatic shifts over the Late Paleogene and Neogene. It is worth noting that, although our divergence ages are generally older than those of most studies, the majority of extant pine diversity is still estimated to have diverged in the Miocene or later. This could point to different drivers being important for the major sectional splits than those that were important for the more recent burst of diversification.

Our study shows that the divergence time estimations depend on the dating method used, as well as the number of fossils and their phylogenetic placement. We cannot judge what method best approximates the pines’ true evolutionary history. Divergence time estimations are dependent on different assumptions inherent in the dating analyses, for example, on the placement of fossils. We urge that future studies relying on dated phylogenetic hypotheses of pines embrace the uncertainty stemming from the different calibration approaches, and that the implicit assumptions between dating approaches are considered. This will increase the robustness and confidence in tested hypotheses and improve our understanding of trait evolutionary processes and its ecological and evolutionary implications.

## Methods

### Taxonomy

We used 115 pine species for phylogenetic inference and followed the taxonomy of Parks *et al.* [6]. We included six additional taxa that were not used in the so far most complete study by Parks *et al.* [6] (*P. balfouriana, P. luchuensis, P. tabuliformis, P. maximinoi, P. tecunumanii, P. jaliscana*).

### DNA sequence matrices

We downloaded eight plastid gene sequences (*matK, rbcL, trnV, ycf1, accD, rpl20, rpoB, rpoC1*) available in GenBank (Additional file 5). We used the sequences provided in Parks *et al*. [6] where possible, supplemented with other sequences from Genbank in a few cases. We ran an automated alignment for all sequences of each gene using MAFFTv7.1 [66], manually checked it and removed ambiguously aligned nucleotides using Gblocks with default settings [67]. The concatenated sequences resulted in a matrix consisting of 115 species and a length of 5866 nucleotides. For 85 species all eight gene sequences were available and for 30 species some sequences were missing (see Additional file 6 for the full sequence matrix also including missing nucleotides).

### Fossil sets

We used two different fossil sets (Additional file 7A). The first set consists of 14 fossils in which we have strong confidence regarding their age and their phylogenetic placement within the genus *Pinus.* The second set of 21 fossils, was more comprehensive and included seven additional fossils that may increase our understanding of divergence times in the genus, although we had somewhat less confidence regarding their exact placement.

We generally selected the fossils according to the following three criteria: (a) the fossil locality could be assigned a precise age, (b) the fossil could be placed to a particular node with high confidence due to specific morphological characters, and (c) the selected fossils are distributed evenly across all major pine clades. In this study, we generally focused on fossils of ovulate cones (except for *P. triphylla*, see Additional file 7A) because other fossil remains (leaves, pollen cones, pollen) are either not commonly preserved or lack the characters relevant to distinguish clades at the subgenus level. The Additional file provides more details on these characters (Additional file 7A), and on the age and placement of each fossil (Additional file 7B-D).

### Phylogenetic reconstruction

We conducted all analyses with BEAST v2.3.1 [68] and constructed the required input-file using BEAUti 2.3.1 [68] with settings described in detail below. We provide all BEAST input files on the Dryad Digital Repository: http://dx.doi.org/10.5061/dryad.74f2r.

#### Partitions, substitution and clock models

We defined the partitions and site models in BEAUti based on the partition scheme and models proposed by PartitionFinder 1.1 [69] (Additional file 8). We applied PartitionFinder using linked branch lengths and the g*reedy* algorithm to search, based on Bayesian Information Criterion (BIC), for the statistically best-fit partitioning schemes and models of nucleotide substitution available in BEAST [68]. Since all gene sequences are from the chloroplast genome and can therefore be expected to be linked, we used the same time-tree for all gene sequences. We further partitioned the clock model and used a separate clock model for the gene sequence *ycf1*, as this gene sequence differed considerably from the others regarding the rate of evolution between lineages. We checked this by comparing the branch lengths of lineages between the single gene trees estimated for each gene sequence separately. For all remaining gene sequences we linked the clock models, as they did not differ much among each other regarding their rate of evolution between lineages. For both clock partitions, we used an uncorrelated relaxed molecular clock model with a log-normal prior.

#### Calibration priors to date the phylogenetic trees

To get estimates for the divergence times in *Pinus*, we used different priors on divergence time to calibrate the molecular clock to an absolute timescale (Additional file 7E). Basically, we applied the traditional node dating method with a birth-death tree prior (ND) and varying node calibration constraints (see details below) and the novel tip dating method with a fossilized birth-death tree prior (FBD) [41, 42], both implemented in BEAST2 [68].

The ND method uses the age of the oldest fossil within a specific clade as a minimum age constraint for the node at which it diverged (calibrating node). We defined these calibrating nodes by determining a monophyletic subset of all the taxa belonging to this clade, so called *taxon sets* (see Additional file 7C/D). For the ND method, a prior calibration density is defined at each calibration node to account for uncertainty underlying the age of the fossil and the likelihood that the true divergence occurred earlier than defined by this fossil record [34]. To compare our analyses with previous studies on pines and to evaluate the sensitivity of ND analyses to prior calibration densities, we used two different approaches to assign prior calibration densities in the ND analyses. In both ND approaches, we used a log-normal distribution for the calibration density at each calibration node, but we varied the shape and breadth of the log-normal distributions.

In the first approach (NDn), we defined a prior calibration density on the calibrating nodes according to the age range of the geological Epoch in which the respective fossil was found. This procedure is commonly used in pines [13, 15]. The offset of the log-normal distribution was set to the minimum age of the corresponding Epoch, whereas the 95^th^ percentile represented the maximum age of the Epoch (Additional file 7E). In the second approach (NDb), we employed a novel procedure for designing the prior calibration density by systematically varying the parameters for the log-normal distribution by fossil age. Specifically, we assumed that the confidence interval (CI) of the priors is narrow for young nodes (5 Ma for the youngest) and higher for the oldest fossils. We increased the CI every 5 Ma by 10% of the previous 5 Ma age class, resulting in a CI of 28 Ma for the oldest (90 Ma) fossil. Hence, we fixed the CI for the youngest fossil to 5 Ma and for the oldest fossil to 28 Ma, then linearly scaled the s.d. of the log-normal priors between 1.0 (youngest fossil) and 0.6 (oldest fossil). This procedure leads to higher densities of young ages close to the minimum fossil age in the calibration priors of the youngest fossils (strong skew), while the prior densities for the oldest fossils are less skewed and their CI spans a broader range of ages (Additional file 7E). We ran each of the prior settings for both sets of fossils, although we could not include all of the listed fossils (Additional file 7) in ND because our technique can only use the oldest fossil of a given clade. We also did not include *P.truckeensis, P.riogrande* and *P.weasmaii* in ND analyses, as it is difficult to justify their node placement without credible synapomorphies. The analyses using the small set (NDns and NDbs) included therefore 12 fossils while analyses using the larger set (NDnl and NDbl) included 15 fossils (Additional file 7B).

In contrast to the traditional ND method, the novel FBD method does not require specification of calibration densities on calibrating nodes to infer absolute ages. It rather includes absolute dates (so called *tip dates*) for extant and extinct taxa or a defined range of dates for a fossil in which the MCMC will sample the fossil uniformly. Here, we fixed the ages to absolute dates. We defined the tip dates as the number of years before the present the specific taxon was living (fossil dates were based on minimum ages listed in Additional file 7 d). As FBD does not require placing fossil constraints to nodes, we could use all of the 14 (FBDs) or 21 (FBDl) fossils from the two fossil sets in the FBD approaches (Additional file 7 e). We additionally ran control analyses (FBDs_ctrl and FBDl_ctrl) with exactly the same fossils (12 and 15) as in ND, to allow for a direct comparison between the two methods. By defining monophyletic taxon sets we either forced the placement of the fossils to the base of a clade and therefore to the specific stem branch (as illustrated for the 16 fossils in Additional file 7 c, black circles), or we set it such that the fossil could likely appear everywhere within a certain clade (as illustrated for the 6 fossils, red circles in Additional file 7 c). Further, FBD analyses were based on rho sampling and not conditioned on the root sampling since the fossil *P. yorkshirensis* is placed along the stem of *Pinus* and the root node represents a sampled node.

### Posterior analysis and summarizing trees

For each setting, we ran two independent analyses in BEAST for either 30×10^7^ or 20×10^7^generations (it turned out that 20×10^7^ generations is by far sufficient to reach convergence and therefore we adjusted some follow up runs to save computational time). We then evaluated the convergence and mixing of the MCMC chains in Tracer v1.6 [68], ensured that the multiple runs converged on the same distribution and ascertained that all ESS (effective sample size) values exceeded 200. We further compared the effective prior and posterior distributions of all the parameters to test whether our analyses are prior-sensitive and the data is informative for the MCMC analyses. We then resampled the resulting files of the inferred phylogenetic trees with a frequency of 10^5^ in logCombiner v2.3.1 [68] and a burn-in of 30% (resp. 46% for the 30x10^7^ generation runs), finally leading to 1401 (resp. 1411) subsampled trees. In ND, we summarized the subsampled trees with a maximum clade credibility tree with common ancestor heights as node heights using TreeAnnotator v2.3.1 [68]. Since we did not provide morphological character data for the fossils and are not interested in the placement of single fossils in the FBD analyses, we pruned off all fossil lineages in all subsampled trees using the *Full2ExtantConvertor.jar*, written by Alexandra Gavryushkina [70]. We summarized these pruned, subsampled FBD trees the same way as in ND.

The posterior age estimates of the subsampled phylogenetic trees of all methods are summarized for 19 selected nodes (node *a-s*) that represent the crown nodes of all major sections in *Pinus*. To test if estimated ages across nodes significantly differ between the methods, we standardized the log-transformed age estimates of every node by first subtracting the mean age across all subsampled trees and methods of that node and second by dividing all ages by the standard deviation of node ages across all subsampled trees and methods of the same node. This yields overall estimated node-ages across trees and methods with a mean of 0 and a standard deviation of 1 for each node. The resulting age differences are now directly comparable across all nodes and allow for estimating the general node-age differences from the overall mean depending on the choice of method and setting (represented as standardized effect size). For this analysis, we used a linear mixed effect model (MCMCglmm [71]) with tree identity as a random effect to account for the inter-dependence of nodes within each of the subsampled posterior trees.

### Prior sensitivity

Priors of multiple calibration constraints can interact and may lead to joint effects, especially when one constraint is ancestral to the other [53], a major shortcoming of ND. We therefore tested if our initially set priors are similar to the effective priors. For this we ran all MCMC analyses without any sequence data to sample only from the prior distribution as recommended by [38], and results illustrated in Additional file 4. In addition, we compared the effective prior calibration densities with the posterior calibration densities to examine the relative influence of the prior and the sequence data on the age estimates [38]. We illustrated this comparison in a figure by plotting the prior against the posterior distribution for both the two ND approaches and the FBD approaches for the small and the large fossil set. For FBD, we illustrated the most recent common ancestral node of the clade the respective fossil was assigned to. We tested whether the methods are significantly more or less sensitive to the priors on time by applying a paired Wilcox test.

### Sensitivity of calibration approaches to single fossil exclusion

To test whether the results are sensitive to the removal of individual fossils we analyzed to what degree the age estimates of the 19 major nodes change in response to removing one single calibration constraint at a time. To do so, we first sampled the 19 node ages from each of the subsampled trees within each analysis when all fossils were used. Next, we ran specific analyses for each method and fossil set, by iteratively leaving out fossils as a calibration constraint, one at a time. Finally, we illustrated the differences in node ages in response to keeping versus removing one single calibration constraint at a time (see Additional file 3 for ND). Note, for the FBD method the most recent common ancestor of the clade a fossil belongs to is represented. These analyses were carried out for both fossil sets.

### Statistical analyses

All statistical analyses and illustrations were generated in the statistical computing environment *R*[72] using the packages phyloch [73], ape [74], geiger [75], raster [76], and MCMCglmm [71].

## Availability of supporting data

The data sets supporting the results of this article are available from the Dryad Digital Repository: http://dx.doi.org/10.5061/dryad.74f2r.

## Authors' contributions

BS, ROW, GL, EC, NS and NEZ designed the study. ABL provided paleontological data for calibration and evaluated potential fossil calibration points. BS performed the phylogenetic analyses, estimated divergence times, analyzed the data and drafted the manuscript. All authors read, provided revisions and approved the manuscript.

## Acknowledgments

BS, NS and NEZ acknowledge support from the Swiss National Science Foundation (grant 31003A_149508/1). We also thank Wilfried Thuiller, Sébastien Lavergne and Tanja Stadler for helpful discussions.

## Additional files

**Additional file 1**: Additional maximum clade credibility (MCC) trees. The trees originate from the node dating (ND) method with narrow prior calibration densities (NDn), with broad prior calibration densities (NDb), and from the fossilized birth-death (FBD) method. Each of these three methods was used in combination of either a small (s) or a large (l) fossil set. The following MCC trees are shown: (A) NDns, (B) NDnl, (C) NDbs, (D) NDbl, (E) FBDs. Note that the MCC tree for FBDl is given in Figure 2. Nodes with red dots indicate Bayesian posterior probabilities lower than 0.95, while all other nodes have posterior probabilities higher than 0.95. Light blue lines on nodes represent the 95% highest posterior density (HPD) of the inferred phylogenetic trees. The node labels (*a-s*) indicate those nodes represented in Figure 3. The geological timescale is in million years. (PDF 210 kb).

**Additional file 2**: Comparison of estimated node ages of the 19 major clades of *Pinus* across all applied dating approaches. **A**: The density of effect sizes originate from a mixed-effect model and illustrate to what degree the estimated node ages differ among dating approaches (different colors; see legend) and among fossil sets (1. darker colors for the large, 2. brighter colors for the small fossil set; see legend). The 95% confidence intervals of effect sizes are illustrated with a line below the density curves. Non-overlap of these intervals indicates significant difference on node ages among all 19 nodes. **B**: Boxplots illustrate the estimated node ages across dating approaches and fossil sets for the major clades (*a-s* illustrated in Figure 2). Whiskers span the 95% highest probability density (HPD), while boxes span the 50% HPD, with the median node age indicated by a vertical bar. The x-axis indicates the geological time in million years. The following abbreviations are used. FBD: fossilized birth-death method; ND: node dating method; l: analyses based on the large fossil set; s: analyses based on the small fossil set; n: narrow calibration priors in ND based on the geological age of the respective fossil; b: broad calibration priors in ND. (PDF 118 kb).

**Additional file 3**: Sensitivity of the time calibration to single fossil exclusion for the node dating approaches (ND). This test measures the difference in age estimates of the 19 major nodes (*a-s*, see also Figure 2) when keeping versus removing single calibration constraints (fossil, labeled on x-axis) at a time. NDns and NDnl are based on narrow prior calibration densities using the small (**A**) and the large (**B**) fossil set, respectively. NDbs and NDbl are based on broad prior calibration densities using the small (**C**) and the large (**D**) fossil set, respectively. Letters (see figure 2 for assignment) indicate nodes with highest deviations. (PDF 126 kb).

**Additional file 4**: Comparison of the specified calibration prior in BEAUti (log-transformed) against the effective calibration prior (log-transformed), as estimated without sequence data, and illustrated for all node dating approaches (ND). NDns and NDnl are based on narrow prior calibration densities using the small (**A**) and the large (**B**) fossil set, respectively. NDbs and NDbl are based on broad prior calibration densities using the small (**C**) and the large (**D**) fossil set, respectively. Labels are only given for fossil constraints with high deviance from the 1:1 line. (PDF 76 kb).

**Additional file 5:** Accession numbers of used gene sequences downloaded from GenBank. (PDF 68 kb).

**Additional file 6**: Sequence matrix used for phylogenetic inference. The number of nucleotide base pairs per gene sequence used for each pine species. N in parentheses gives the number of positions in this gene sequence for which the nucleotide pair is undetermined. (PDF 66 kb).

**Additional file 7:** Summary of fossil information. **A:** the single fossils used for calibration constraints are first listed and then discussed, ordered by taxonomic group. In the section where fossils are discussed, bold-italic font represent those fossils used in the smaller fossil set, while italic font represents those used in the larger fossil set only. The listed age indicates the minimum age of the corresponding fossil. **B:** Summary information of fossils used. The character “x” indicates in which dating approach the single fossil was used. The geological layer (and the time scale thereof in Ma) lists, were the fossil was excavated. The reference indicates the publication from which the description of the fossil was taken. Asterisks on fossil numbers represent those used in the large fossil set only. Abbreviations used are FBD: fossilized birth-death method; ND: node dating method; l: denotes analyses based on the large fossil set; s: denotes analyses based on the small fossil set; n: narrow calibration priors in ND based on the geological age of the respective fossil; b: broad calibration priors in ND. **C**: Fossil placement for the ND approaches; asterisks on fossil numbers represent those used in the large fossil set only. **D**: Fossil placement for the FBD approaches; asterisks on fossil numbers represent those used in the large fossil set only. Red dots indicate fossils that were allowed to be placed anywhere within the corresponding clade. Black dots illustrate fossils that were allowed to be placed anywhere along the indicated branch. **E**. Specified prior calibration densities used for ND approaches (NDn with narrow, NDb with broad densities) and absolute tip dates of fossils used in the fossilized birth-death approach (FBD) are listed. Asterisks on fossil number represent those used in the large fossil set only. (PDF 282 kb).

**Additional file 8**: Settings and output summary of PartitionFinder showing the best partition scheme for the used sequence data in this study. (PDF 31 kb).

